# Manumycin Polyketides Act as Molecular Glues Between UBR7 and P53 to Impair Breast Cancer Pathogenicity

**DOI:** 10.1101/814285

**Authors:** Yosuke Isobe, Mikiko Okumura, Ross White, Lynn M. McGregor, Jeffrey M. McKenna, John A. Tallarico, Markus Schirle, Thomas J. Maimone, Daniel K. Nomura

**Affiliations:** Department of Chemistry, University of California, Berkeley, Berkeley, CA 94720 USA; Novartis-Berkeley Center for Proteomics and Chemistry Technologies; Novartis Institutes for BioMedical Research, Cambridge, MA 02139 USA; Department of Molecular and Cell Biology, University of California, Berkeley, Berkeley, CA 94720 USA; Department of Nutritional Sciences and Toxicology, University of California, Berkeley, Berkeley, CA 94720 USA; Innovative Genomics Institute, Berkeley, CA 94704 USA

**Keywords:** manumycin, asukamycin, molecular glues, E3 ligases, chemoproteomics, activity-based protein profiling, natural products, covalent ligands, undruggable, protein-protein interaction (PPI)

## Abstract

Molecular glues are an intriguing therapeutic modality that harness small-molecules to induce interactions between proteins that typically do not interact, thus enabling the creation of novel protein functions not naturally encoded in biology. While molecular glues such as thalidomide and rapamycin have catalyzed drug discovery efforts, such molecules are rare and have often been discovered fortuitously, thus limiting their potential as a general strategy for therapeutic intervention of disease. Historically, natural products have proven to be important sources of molecular glues and we postulated that natural products bearing multiple electrophilic sites may be an unexplored source of such molecules, potentially through multi-covalent attachment. Using activity-based protein profiling (ABPP)-based chemoproteomic platforms, we show that members of the manumycin family of polyketides, which bear multiple potentially reactive sites, target C374 of the putative E3 ligase UBR7 in breast cancer cells to impair breast cancer pathogenicity through engaging in molecular glue interactions with the neo-substrate tumor-suppressor TP53, leading to the activation of p53 transcriptional activity and cell death. Our results reveal a previously undiscovered anti-cancer mechanism of this natural product family and highlight the potential for combining chemoproteomics and multi-covalent natural products for the discovery and characterization of new molecular glues.

## Introduction

One of the biggest challenges faced in discovering new cancer therapies is that many genetically validated targets are considered “undruggable,” in that they do not possess known binding pockets or “druggable hotspots” that can be functionally targeted with small-molecules using traditional drug discovery paradigms ^1^. The grand challenge of tackling the undruggable proteome has inspired the development of innovative technologies that enable novel approaches for functional targeting biomolecules with new therapeutic modalities. Examples of some of these small molecule-based technologies include chemoproteomics-enabled covalent ligand screening, DNA-encoded library platforms for discovering ligands against undruggable proteins, proteolysis-targeting chimeras (PROTACs) or degraders for targeted ubiquitin-proteasome system-dependent degradation of proteins, and “molecular glues” for small-molecule induced formation of protein interfaces that confer enhanced, inhibited, or new protein function ^2–9^. Molecular glue-based degraders, as exemplified by the IMiD-family of immunomodulatory drugs including thalidomide, are notable for their potentially lower molecular weight as compared to linker-based bifunctional molecules (e.g. PROTACS), a possible advantage for increased bioavailability and improved pharmacokinetic profiles ^7^. Moreover, molecular glues are also particularly interesting from a functional perspective, since the few well-characterized examples uniquely modulate protein function and downstream biology by stabilizing of complexes between protein binding partners that otherwise would not interact.

Three notable, well-characterized molecular glues showcase diverse functional outcomes associated with novel protein complexes. The natural product rapamycin binds to FKBP12, leading to the recruitment of mTORC1, and resulting in partial inhibition of mTORC1 signaling, which exerts multiple therapeutic effects.^10^ Fusicoccin and cotylenin diterpenes form interfaces between the 14-3-3 protein family and a variety of interacting partners to modulate signal transduction, apoptosis and cell-cycle control, and cell differentiation ^11,12^. Thalidomide binds to the E3 ligase cereblon to form a new protein interface that recruits neo-substrates such as SALL4 or Ikaros transcription factors for ubiquitination and degradation ^13–16^. Paradoxically, degradation of the former leads to teratogenic effects while degradation of the latter is a proven chemotherapeutic strategy^7,9,17^. Moreover, analogs of rapamycin and thalidomide can maintain their respective binding to their FKBP12 and cereblon targets but shift their neo-substrate binding profiles to confer new and distinct biological functions ^18–20^. This realization has led to the discovery of approved drugs such as lenalidomide, pomalidomide, FK506. Unfortunately, only a handful of well-characterized molecular glues have been discovered; thus, limiting our understanding of the design principles behind their function and reducing their impact on drug discovery. Natural products have been a robust source of many known molecular glue interactions (e.g. rapamycin, FK506, auxin, brefeldin A, forskolin, and others), and their diverse and stereochemically-rich architectures possess many positive attributes for recruiting protein surfaces ^9,17^.

Electrophilic natural products represent a vastly underexplored subset of natural products from which molecular glues can originate, and such species could have unique and positive attributes owing to their potential for covalent bond formation^21^. While covalently-acting natural products have not been previously identified as molecular glues, we hypothesized that natural products that bear: 1) more than one potential site for covalent interaction with nucleophilic amino acids on proteins; and 2) have also been shown to possess biological activity, would constitute candidates for possessing potentially unique glue-like characteristics.

The covalency of these potential natural product-based molecular glues also enables rapid and facile mechanistic deconvolution of direct protein targets using chemoproteomic platforms. One such chemoproteomic platform used herein is activity-based protein profiling (ABPP), which uses reactivity-based chemical probes to profile proteome-wide reactive, functional, and ligandable hotspots directly in complex living systems. When used in a competitive manner, covalently-acting small-molecules can be competed against the binding of reactivity-based probes to facilitate the identification of their protein targets and covalent sites of modification 3,4,22–24. Upon identifying the direct targets of the natural product, standard pull-down and quantitative proteomic profiling experiments can be used to identify specific molecular glue interactions enabled by the natural product.

In this study, we interrogated the *Streptomyces-*derived family of manumycin polyketide natural products, specifically asukamycin and manumycin A, which have documented antibiotic and antiproliferative properties ^25^. Manumycin A has also been show to inhibit farnesyl transferase activity and sphingomyelinase activity, but asukamycin is not known to inhibit these targets and its anti-cancer mechanism of action is unclear ^26–31^. These molecules possess a suite of electrophilic sites that could potentially react covalently with nucleophilic amino acids within proteins, such as cysteines **(Figure 1a)** ^26^. We hypothesized that multiple reactive sites within these natural products may lead to specific interactions with multiple protein targets resulting in bifunctional or molecular glue-type interactions.

**Figure 1.**
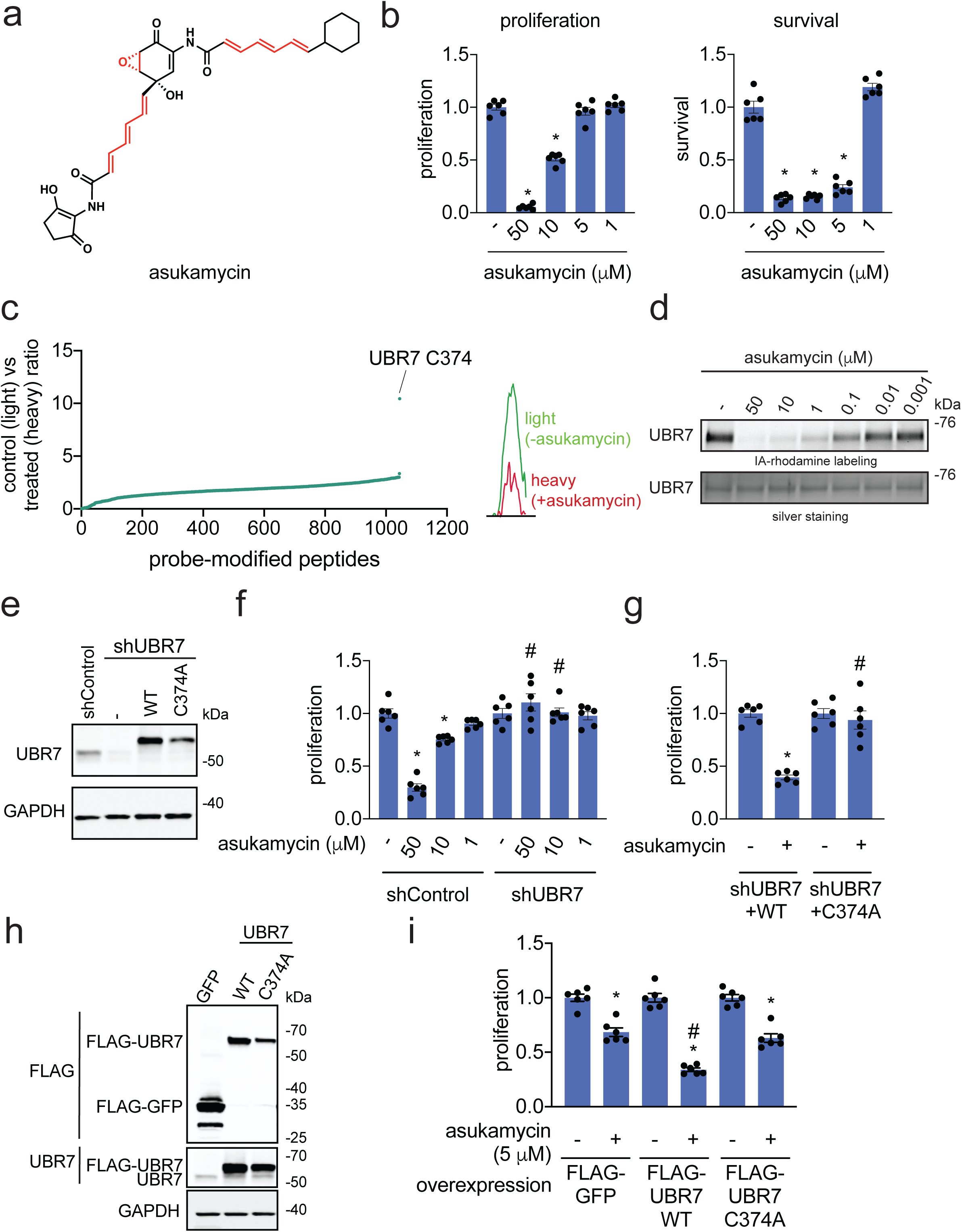
Asukamycin targets C374 of UBR7 to impair breast cancer cell proliferation. **(a)** Structure of asukamycin highlighting (in red) three potential reactive sites. **(b)** Proliferation and serum-free cell survival in 231MFP breast cancer cells treated with DMSO vehicle or asukamycin for 48 h, assessed by Hoechst stain. **(c)** IsoTOP-ABPP analysis of asukamycin *in situ* in 231MFP cells. 231MFP cells were treated with DMSO vehicle or asukamycin (10 μM) for 3 h, and resulting cell lysates were labeled with IA-alkyne (100 μM) for 1 h after which isotopically light (control) or heavy (asukamycin-treated)-treated TEV tag-bearing biotin-azide handles were appended by CuAAC and taken through the isoTOP-ABPP method. Shown are light to heavy ratios of probe-modified peptides. Shown are average probe-modified peptide ratios. The individual replicate values and total datasets are shown in **Supplementary Table 1**. On the right is a representative MS1 chromatogram of the probe-modified C374-bearing UBR7 peptide between control versus asukamycin-treated cells. **(d)** Gel-based ABPP analysis of asukamycin against pure human UBR7 protein. UBR7 protein was pre-incubated with DMSO vehicle or asukamycin for 30 min prior to IA-rhodamine labeling of UBR7 for 1 h at room temperature. Protein was resolved by SDS/PAGE and visualized by in-gel fluorescence and protein loading was assessed by silver staining. **(e)** Expression of UBR7 and loading control GAPDH levels in 231MFP shControl, shUBR7, or shUBR7 cells expressing FLAG-tagged wild-type (WT) or FLAG-tagged C374A mutant UBR7 protein, assessed by Western blotting. **(f)** Cell proliferation in 231MFP shControl or shUBR7 cells treated with DMSO vehicle or asukamycin for 24 h, assessed by Hoechst stain. **(g)** Cell proliferation in 231MFP shUBR7 cells expressing FLAG-WT or FLAG-C374A mutant UBR7 treated with DMSO vehicle or asukamycin (50 μM) for 24 h. **(h)** anti-FLAG, anti-UBR7, and loading control anti-GAPDH protein expression in 231MFP cells stably expressing FLAG-GFP, FLAG-UBR7 WT, or FLAG-UBR7 C374A mutant assessed by Western blotting. **(i)** Cell proliferation of 231MFP cells stably expressing FLAG-GFP, FLAG-UBR7 WT or FLAG-UBR7 C374A mutant treated with DMSO or asukamycin (5 μM) for 48 h, assessed by Hoechst stain. Data shown in (**b, f, g**, and **i)** are shown as individual replicate values and average ± sem and are n=6 biologically independent samples/group. Gels shown in **(d, e**, and **h)** are representative blots from n=3 biologically independent samples/group. Statistical significance was calculated with two-tailed unpaired Student’s t-tests and are shown as *p<0.05 compared to vehicle-treated controls within each group, #p<0.05 against respective concentrations of asukamycin-treatment in shControl groups in **(f, g)** or asukamycin-treatment in FLAG-GFP expressing groups in **(i)**.

Using ABPP-based chemoproteomic platforms, we demonstrated that one of the primary targets of asukamycin is C374 of the postulated E3 ubiquitin ligase UBR7. We further showed that this UBR7-asukamycin complex engages multiple proteins in human triple negative breast cancer cells including the tumor suppressor TP53, and that the resulting UBR7-asukamycin-TP53 complex contributes to the anti-cancer activity of asukamycin.

## Results

### Anti-Cancer Activity of Asukamycin

There is significant unmet medical need for new triple-negative breast cancer (TNBC) therapies as such malignancies have worse clinical prognoses than other breast cancer subtypes. Moreover, few targeted therapies are approved for the treatment of TNBCs. New small-molecules, mechanisms, and therapeutic modalities for combatting TNBCs could greatly reduce mortalities associated with these aggressive breast cancers ^32^. We show that asukamycin impairs the cell survival and proliferation of two TNBC-derived cell lines, 231MFP and HCC38 **(Figure 1b, Supplementary Figure 1)**.

### ABPP to Map Asukamycin Targets

We postulated that the antiproliferative activity of asukamycin may be dependent on covalent interactions with cysteines on specific protein targets. Asukamycin could react with protein nucleophiles through either hetero-Michael addition reactions with one or both of the two polyunsaturated amide side chains or reaction with the epoxyketone moiety, a known process for similar epoxy ketone natural products. ^33,34^ We performed ABPP-based chemoproteomic profiling to identify asukamycin targets in 231MFP TNBC cells. Proteome samples treated with vehicle or asukamycin were labeled with the cysteine-reactive probe iodoacetamide-alkyne (IA-alkyne or *N-*hex-5-ynyl-2-iodo-acetamide), prior to further processing using well-validated methods for covalent ligand target identification ^4,22–24,35,36^. We identified C374 of UBR7 as a primary target of asukamycin with the highest control to treated (or light to heavy) ratio in 231MFP cells **(Figure 1c, Supplementary Dataset 1)**. We confirmed that UBR7 was a direct target of asukamycin using gel-based ABPP approaches, wherein we show competition of asukamycin against rhodamine-functionalized iodoacetamide (IA-rhodamine) labeling of recombinant human UBR7 protein **(Figure 1d)**.

Next, we investigated whether the anti-cancer effects of asukamycin were driven through asukamycin interactions with C374 of UBR7. Stable UBR7 knockdown in 231MFP cells did not compromise proliferation but conferred complete resistance to asukamycin-mediated anti-proliferative effects **(Figure 1e, 1f)**. Expression of wild-type (WT) or C374A mutant UBR7 in shUBR7 231MFP cells led to re-sensitization of asukamycin-mediated anti-proliferative effects in WT but not C374A UBR7-expressing cells **(Figure 1g)**. Taken together, these results demonstrate that asukamycin targets C374 of UBR7 and that this interaction is responsible for the anti-proliferative effects of asukamycin in breast cancer cells. These data also hinted that asukamycin, through targeting C374 of UBR7, may be conferring a gain-of-function effect, since UBR7 knockdown in itself does not impair cell proliferation **(Supplementary Figure 2)**. Further supporting this hypothesis, we found that overexpression of WT, but not C374A mutant FLAG-UBR7 conferred significantly heightened asukamycin-induced anti-proliferative effects **(Figure 1h, 1i)**.

While UBR7 is annotated as an E3 ubiquitin ligase and postulated to be involved in histone ubiquitination 37, its biochemical and physiological functions are poorly understood especially with regard to cancer cell proliferation. We attempted to reconstitute the activity of UBR7 *in vitro*, but our various efforts to demonstrate UBR7 activity failed. Instead, we next explored the potential for UBR7 to engage in asukamycin-induced interactions with other proteins.

### Mapping Molecular Glue Interactions of UBR7-Asukamycin

From these results and the multi-covalent potential of asukamycin, we postulated that asukamycin may be conferring gain-of-function effects upon UBR7 through engaging in molecular glue interactions with other proteins. To substantiate this hypothesis, we performed proteomic analysis on anti-FLAG pulldown eluate from FLAG-GFP or FLAG-UBR7-expressing 231MFP cells treated with vehicle or asukamycin to identify protein-protein interactions that were dependent on both UBR7 *<u>and</u>* asukamycin. From this experiment, we identified 13 proteins that showed significant (p<0.01) and >15-fold higher pulldown in the asukamycin-treated Flag-UBR7 cells compared to vehicle-treated FLAG-UBR7 cells **(Figure 2a, Supplementary Dataset 2)**. Among these 13 targets, 8 are UBR7-dependent, with >4-fold enrichment between asukamycin-treated FLAG-UBR7 cells versus asukamycin-treated FLAG-GFP cells **(Figure 2a, Supplementary Dataset 2)**. These 8 targets include proteins of great importance in cancer, including DNA protein kinase (PRKDC) and the tumor-suppressor p53 (TP53) ^38–40^ **(Figure 2b)**. In pulldown experiments, we confirmed the asukamycin-dependent interactions with UBR7 with Western blots showing that proteins such as TP53 and PRKDC interacted specifically in FLAG-UBR7 expressing cells treated with asukamycin **(Figure 2c)**. Interestingly, in our anti-FLAG blot from FLAG-UBR7 pulldown studies, we also observed several higher molecular weight species that corresponded to the estimated combined masses of FLAG-UBR7 and TP53 as well as even higher molecular weight species that corresponded to the estimated molecular weight of PRKDC. In the anti-TP53 blot from the same pulldown studies, in addition to the expected TP53 band, we also observed a distinct higher molecular weight species that corresponded to a molecular weight in line with the addition of FLAG-UBR7 and TP53 **(Figure 2c)**. Reinforcing that this higher molecular weight FLAG-UBR7 band included the TP53 higher molecular weight species, a dual color Western blot showed overlap between the FLAG-UBR7 and TP53 higher molecular weight bands **(Supplementary Figure 3)**. UBR7 knockdown in 231MFP cells treated with asukamycin also resulted in >80 % reduction in the higher molecular weight TP53 species compared to shControl counterparts **(Figure 2d, Supplementary Figure 4)**. TP53 knockdown in 231MFP cells treated with asukamycin resulted in a significant, albeit less pronounced reduction in the higher molecular weight FLAG-UBR7 species compared to shControl counterparts **(Supplementary Figure 4)**. While there may be additional proteins of similar molecular weight to TP53 that are included in this higher molecular weight FLAG-UBR7 species, we believe that TP53 is one of those proteins. We did not observe higher molecular weight PRKDC bands likely because of the poor resolution of proteins at this high molecular weight. We further confirmed that these higher molecular weight species were not polyubiquitinated TP53 since treatment of proteomes with the deubiquitinase USP2, which eliminated total proteome ubiquitination, was not able to eliminate these higher molecular weight TP53 bands **(Supplementary Figure 5)**. These results show enrichment of both the parent molecular weight protein interaction partner and also a higher molecular weight species, suggesting that the complexes between UBR7-asukamycin and binding partners such as TP53 or PRKDC initially form through reversible interactions which eventually in-part lead to multi-covalent interactions. We also showed that abundant proteins such as GAPDH did not interact with the asukamycin-UBR7 complex, arguing for specific molecular interactions over non-specific binding events **(Figure 2c)**. While TP53 was not identified as a target in our isoTOP-ABPP studies likely due to its low abundance or the potential interaction of asukamycin with another nucleophilic amino acid besides cysteine, we further demonstrated that asukamycin directly interacts with TP53 by gel-based ABPP **(Figure 2e)**.

**Figure 2.**
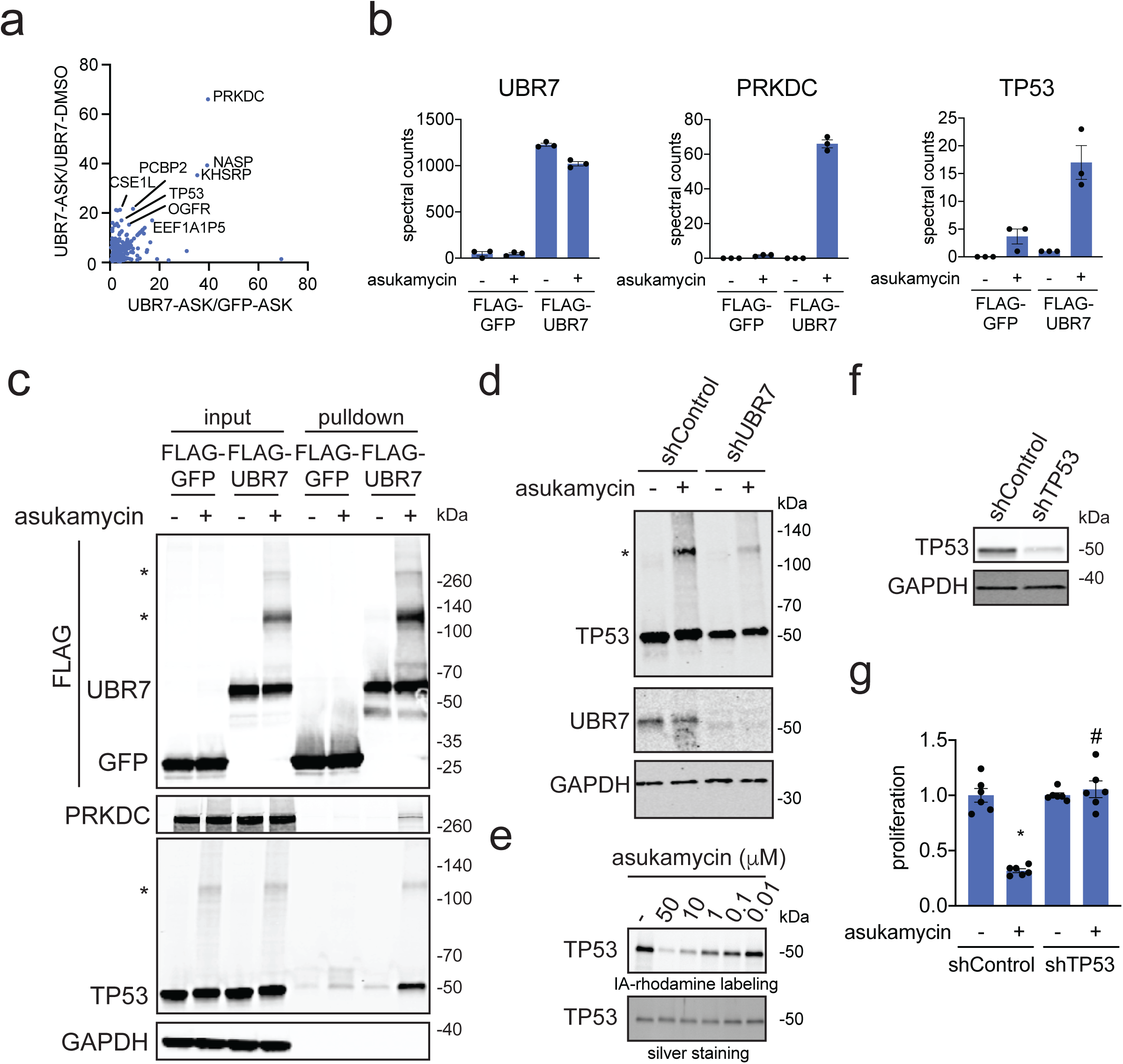
Mapping molecular glue interactions of asukamycin and UBR7. **(a)** Proteomics analysis of molecular glue interactions of UBR7-asukamycin. 231MFP cells stably expressing FLAG-GFP or FLAG-UBR7 were treated with DMSO or asukamycin (50 μM) for 3 h. FLAG-GFP and FLAG-UBR7 interacting proteins were subsequently enriched and then subjected to proteomic analysis. Raw and analyzed proteomic analysis is shown in **Supplementary Table 2** and is quantified by spectral counting. For those proteins that showed no peptides in a particular sample group, we set those proteins to 1 to enable relative fold-change quantification to generate this figure to visually show those proteins that showed high enrichment in asukamycin (ASK)-treated FLAG-UBR7 groups compared to asukamycin-treated FLAG-GFP groups on the x-axis and ASK-treated FLAG-UBR7 groups compared to DMSO-treated FLAG-UBR7 groups. Proteins highlighted showed >15-fold enrichment comparing UBR-ASK to UBR7-DMSO groups and >4-fold enrichment comparing UBR7-ASK to GFP-ASK groups. **(b)** Shown are spectral counts for representative proteins in the experiment described in **(a). (c)** Protein levels of FLAG-tagged proteins, PRKDC, TP53, and loading control GAPDH from a repeat of the experiment described in **(a)** by Western blotting showing specific interactions of PRKDC and TP53 with FLAG-UBR7 only when cells are treated with asukamycin. Higher molecular weight species of FLAG-UBR7 and TP53 are noted with (*). **(d)** anti-TP53, anti-UBR7, and loading control anti-GAPDH Western blots in shControl or shUBR7 231MFP cells treated with vehicle DMSO or asukamycin (50 μM) for 3 h. Higher molecular species of TP53 is noted with (*). Quantification for this gel is provided in **Supplementary Figure 4a. (e)** Gel-based ABPP analysis of asukamycin against pure human TP53 protein. TP53 protein was pre-incubated with DMSO vehicle or asukamycin for 30 min prior to IA-rhodamine labeling of TP53 for 1 h at room temperature. Protein was resolved by SDS/PAGE and visualized by in-gel fluorescence and protein loading was assessed by silver staining. **(f)** TP53 protein levels in 231MFP shControl and shTP53 cells. **(g)** Cell proliferation of 231MFP shControl and shTP53 cells treated with DMSO or asukamycin (10 μM) for 24 h. Data shown in **(b** and **g)** are shown as individual replicate values and average ± sem and are n=3 for **(b)** and n=6 for **(g)** biologically independent samples/group. Gels shown in **(c, d, e**, and **f)** are representative blots from n=3 biologically independent samples/group. Statistical significance was calculated with two-tailed unpaired Student’s t-tests and are shown as *p<0.05 compared to vehicle-treated controls within each group, #p<0.05 compared to asukamycin-treated shControl group in **(g)**.

Given the importance of PRKDC and TP53 in cancer pathogenicity, we next tested the relative contributions of these two proteins in asukamycin-mediated anti-proliferative effects. TP53 knockdown, but not PRKDC knockdown, conferred complete resistance to asukamycin-mediated anti-proliferative effects in 231MFP breast cancer cells, indicating that TP53 is the more critical target **(Figure 2f, 2g, Supplementary Figure 6)**. Taken together, these results suggest that asukamycin has the ability to promote the formation of a gain-of-function complex between UBR7 and TP53 that results in anti-proliferative effects in TNBC cell lines.

### Characterizing UBR7-Asukamycin Molecular Glue Interactions with TP53

We next investigated the biochemical and functional consequences of UBR7, asukamycin, and TP53 interactions. We wondered whether the UBR7-asukamycin-TP53 complex might affect TP53 function in the TP53-mutant setting of TNBC cell lines. Consistent with asukamycin directly binding to TP53 and potentially stabilizing TP53 folding, asukamycin caused a significant increase in TP53 thermal stability *in vitro* in 231MFP cell lysate **(Figure 3a, 3b)**. We also observed a higher molecular weight TP53 species, which showed even greater thermal stability compared to the asukamycin-treated TP53 parental protein **(Figure 3a, 3b)**. Asukamycin also significantly increased TP53 binding to its DNA consensus sequence *in vitro* when TP53 was spiked into 231MFP cell lysate, indicating that asukamycin potentially activated TP53 transcriptional activity **(Figure 3c)**. As other thiol-reactive compounds have been reported to both thermally stabilize TP53 and to rescue the activity of mutant TP53 ^41^, we next asked if asukamycin could increase the transcriptional activity of TP53. Consistent with functional activation of p53 activity in cells, asukamycin induced the expression of the p53 target gene TP53AIP1 in 231MFP cells. The asukamycin-mediated increase in expression was attenuated in UBR7 knockdown cells **(Figure 3d)**. Asukamycin also significantly induced p53 luciferase reporter transcriptional activity in HEK293T cells, even more so than the positive control topoisomerase inhibitor doxorubicin (dox) that is known to induce apoptosis through activating the p53 pathway **(Figure 3e)**. This asukamycin-induction of p53 transcriptional reporter activity was completely attenuated in cells expressing C374A mutant UBR7 compared to wild-type UBR7 in shUBR7 HEK293T cells **(Figure 3f, 3g)**. Collectively, these results indicated that asukamycin directly targets TP53, stabilizes TP53 folding, and activates p53 transcriptional activity in a UBR7 C374-dependent manner.

**Figure 3.**
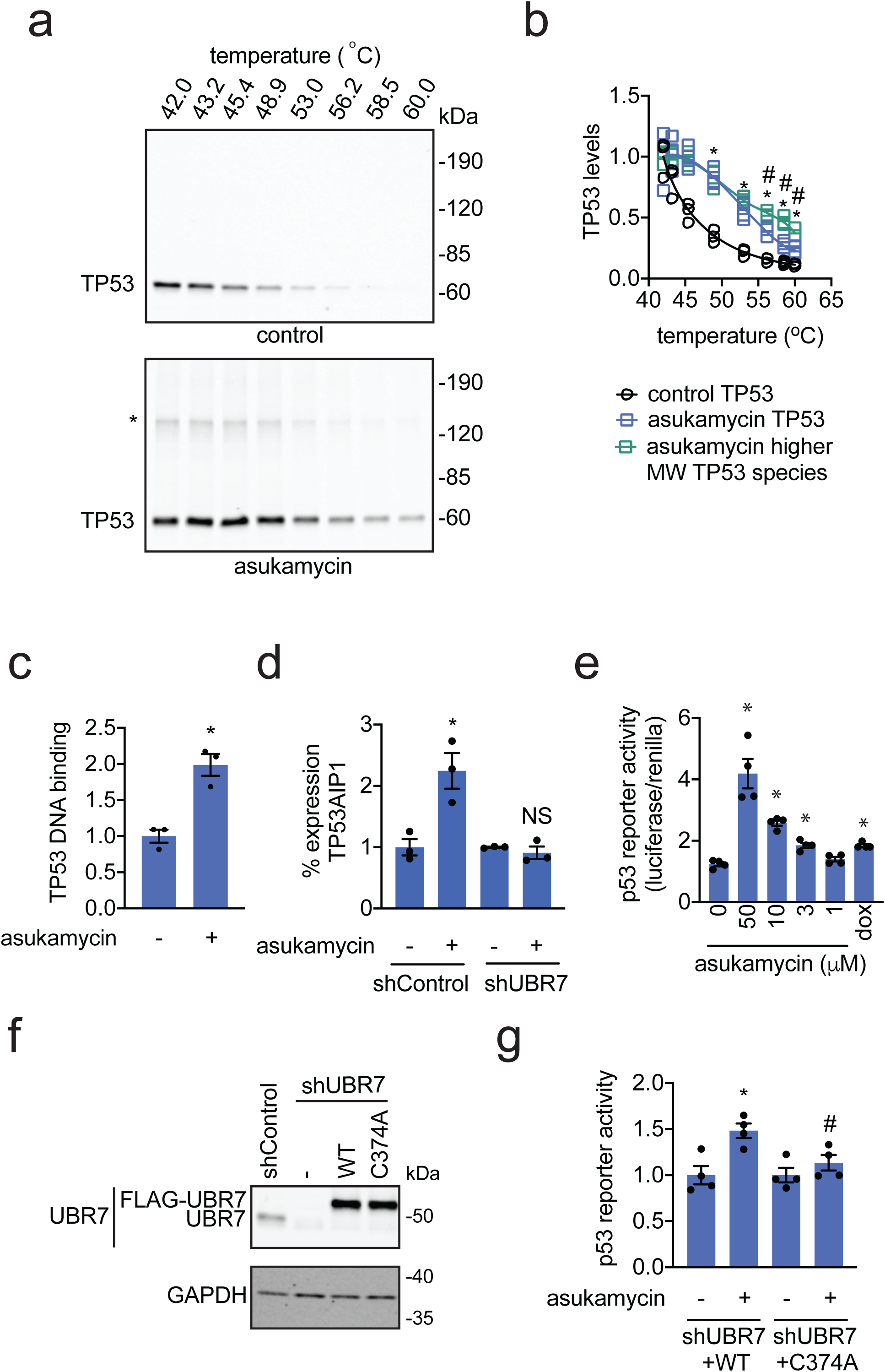
Asukamycin acts as a molecular glue between UBR7 and TP53 and activates TP53 transcriptional activity in a UBR7-dependent manner. **(a)** Thermal shift assay *in vitro* in 231MFP breast cancer cell lysate treated with DMSO vehicle or asukamycin (50 μM). Lysate was heated to the designated temperature for 3 minutes, followed by 3 minutes at room temperature, centrifugation, and SDS/PAGE and Western blotting for TP53. **(b)** Quantification of TP53 parent and higher molecular weight levels from thermal shift assay described in **(a)**. Data is normalized to respective 42 °C TP53 protein levels. **(bc)** TP53 DNA binding to p53 DNA consensus sequence *in vitro* with TP53 spiked into 231MFP breast cancer cell lysate treated with DMSO vehicle or asukamycin (50 μM). **(d)** mRNA expression of TP53AIP1 in 231MFP shControl and shUBR7 cells treated with DMSO vehicle or asukamycin (50 μM) for 6 h, assessed by qPCR. **(e)** p53 reporter activity reported as the ratio between luciferase reporter activity versus cell number control renilla levels in shUBR7 HEK293T cells expressing WT or C374A mutant UBR7 and the p53 reporter construct treated with DMSO vehicle control, asukamycin, or doxorubicin (1 μM) for 6 h. **(f)** Protein expression of UBR7 and loading control GAPDH in HEK293T shControl and shUBR7 cells expressing empty vector, wild-type UBR7, or C374A mutant UBR7 assessed by Western blotting. **(g)** p53 reporter activity reported compared to DMSO vehicle-treated controls in each group in HEK293T shUBR7 cells expressing the p53 reporter construct treated with DMSO vehicle control or asukamycin (50 μM) for 24 h. Data shown in (**b, c, d, e**, and **g)** are shown as individual replicate values and average ± sem and are n=3 for **(b, c**, and **d)** and n=4 for **(e** and **g)** biologically independent samples/group. Gels shown in **(a** and **f)** are representative blots from n=3 biologically independent samples/group. Statistical significance was calculated with two-tailed unpaired Student’s t-tests and are shown as *p<0.05 compared to each temperature control group for **(b)**, vehicle-treated controls for **(c, d, e**, and **g)** within each group, #p<0.05 compared to corresponding asukamycin treatment groups for each temperature for parent TP53 species in **(b)** and asukamycin-treated shUBR7 WT group in **(g)**. NS denotes not significant.

### Characterizing the activity of related natural product manumycin A

Asukamycin belongs to a larger family of polyketide natural products known as manumycins. While manumycin A, a close structural analog of asukamycin that also bears multiple electrophilic sites, has been shown to inhibit farnesyl transferase or sphingomyelinase ^26–31^ **(Figure 4a)**, the effect of manumycin A on UBR7 or its molecular glue activity with TP53 have not been previously reported. Interestingly, manumycin A also bound to UBR7 at comparable potency to asukamycin by gel-based ABPP **(Figure 4b)**. We also found that manumycin A, like asukamycin, activated p53 transcriptional activity in HEK293T cells more so than doxorubicin **(Figure 4c)**. Manumycin A treatment also led to molecular glue interactions between UBR7 and TP53 in 231MFP cells and resulted in distinct higher molecular weight species which correspond to the added molecular weight of TP53 and UBR7 **(Figure 4d)**. Finally, manumycin A, like asukamycin, interacted directly with TP53 by gel-based ABPP **(Figure 4e)**.

**Figure 4.**
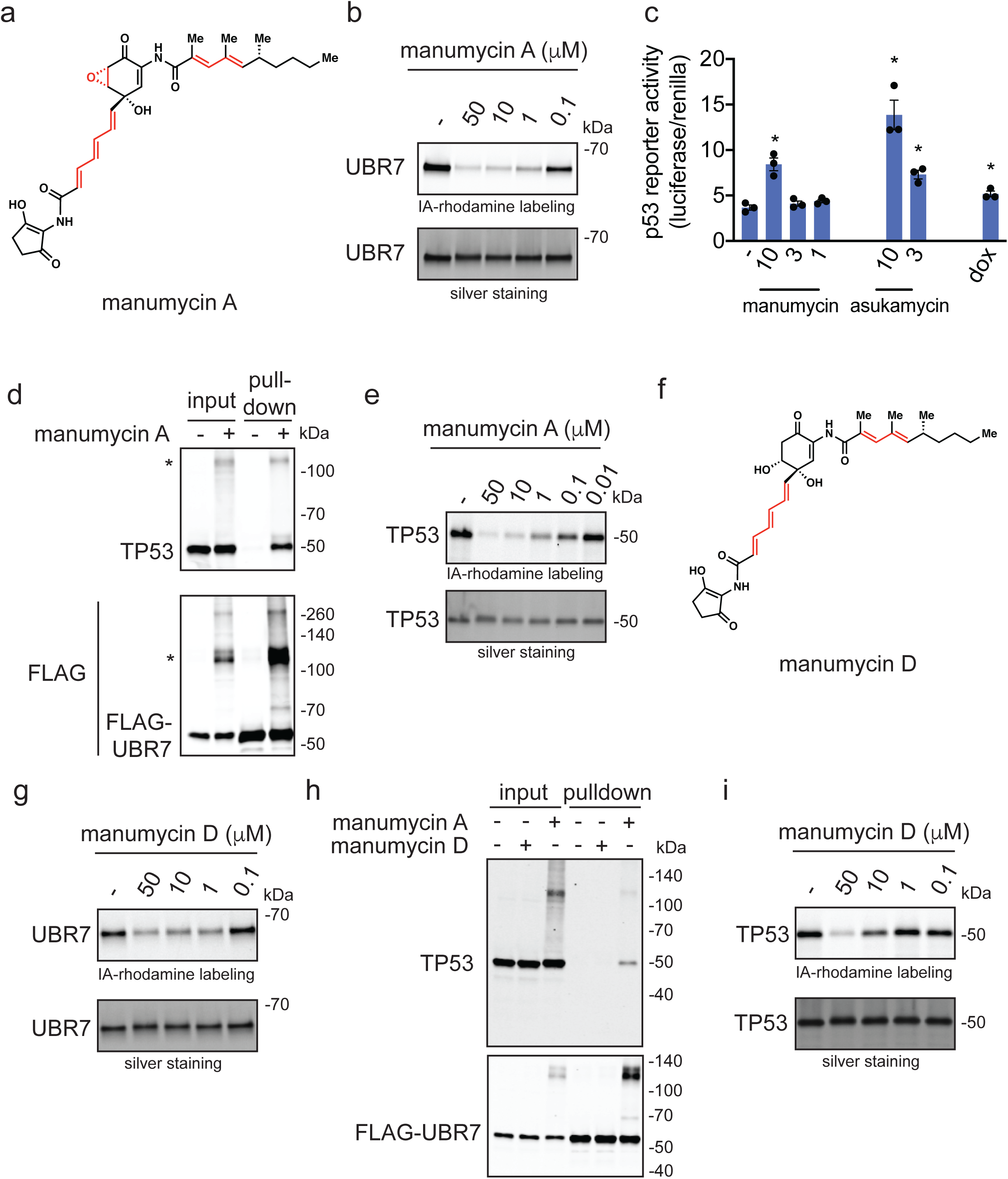
Manumycin A also interacts with UBR7 and engages in molecular glue activities with TP53. **(a)** Structure of manumycin A highlighting the potential reactive sites in red. **(b)** Gel-based ABPP analysis of manumycin A against pure human UBR7 protein. UBR7 protein was pre-incubated with DMSO vehicle or manumycin A for 30 min prior to IA-rhodamine labeling of UBR7 for 1 h at room temperature. Protein was resolved by SDS/PAGE and visualized by in-gel fluorescence and protein loading was assessed by silver staining. **(c)** p53 reporter activity reported as the ratio between luciferase reporter activity versus cell number control renilla levels in HEK293T cells expressing the p53 reporter construct treated with DMSO vehicle control, manumycin A, asukamycin, or doxorubicin (1 μM) for 6 h. **(d)** Protein levels of TP53 and FLAG-tagged proteins by Western blotting in 231MFP cells stably expressing FLAG-UBR7 treated with DMSO vehicle or manumycin A (50 μM) for 3 h, after which FLAG-UBR7 interacting proteins were subsequently enriched. Higher molecular weight TP53 and FLAG-UBR7 species are noted with a (*). **(e)** Gel-based ABPP analysis of manumycin A against pure human TP53 protein. TP53 protein was pre-incubated with DMSO vehicle or manumycin A for 30 min prior to IA-rhodamine labeling of TP53 for 1 h at room temperature. **(f)** Structure of manumycin D with reactive sites highlighted in red. **(g)** Gel-based ABPP analysis of manumycin D against pure human UBR7 protein. UBR7 protein was pre-incubated with DMSO vehicle or manumycin D for 30 min prior to IA-rhodamine labeling of UBR7 for 1 h at room temperature. **(h)** Protein levels of TP53 and FLAG-tagged proteins by Western blotting in 231MFP cells stably expressing FLAG-UBR7 treated with DMSO vehicle or manumycin D (50 μM) for 3 h, after which FLAG-UBR7 interacting proteins were subsequently enriched. **(i)** Gel-based ABPP analysis of manumycin D against pure human TP53 protein. TP53 protein was pre-incubated with DMSO vehicle or manumycin D for 30 min prior to IA-rhodamine labeling of TP53 for 1 h at room temperature. Data shown in (**c)** are shown as individual replicate values and average ± sem and are n=3 biologically independent samples/group. Gels shown in **(b, d, e, g, h**, and **i)** are representative blots from n=3 biologically independent samples/group. Statistical significance as calculated with two-tailed unpaired Student’s t-tests and are shown as *p<0.05 compared to vehicle-treated controls.

To investigate the relative contribution of the epoxide warhead on UBR7 binding versus TP53 binding, we prepared manumycin D **(Figure 4f)** ^42,43^, an epoxide-reduced variant, in one step from manumycin A (see **Synthetic Methods)** ^44,45^. Quite interestingly, manumycin D still labeled UBR7 by gel-based ABPP, albeit weaker than manumycin A **(Figure 4g)**, but was incapable of inducing interactions between UBR7 and TP53 **(Figure 4h)**. Consistent with these data, we also showed that manumycin D bound less potently to TP53 compared to manumycin A or asukamycin by gel-based ABPP **(Figure 4i)**. Moreover, manumycin A, but not manumycin D, impaired 231MFP breast cancer cell proliferation **(Supplementary Figure 7)**. These data collectively suggest that the epoxide contributes to interactions with TP53 and is necessary to engage in multi-covalent interactions between UBR7 and TP53 and exert anti-proliferative effects. These results also imply that the unsaturated side chains of manumycin A, and by analogy asukamycin, were responsible for the covalent interaction with C374 of UBR7.

## Discussion

Molecular glues are intriguing druggable modalities that harness small-molecules to confer new and interesting properties to proteins for potential therapeutic applications. While this drug discovery modality has the potential for stabilizing interactions between proteins that usually would not interact and thus enables exploitation of unique protein functions, discovery of new molecular glue scaffolds have been mostly serendipitous. We postulated that natural products bearing multiple electrophilic sites may engage in multi-covalent interactions that bring specific proteins into novel, functionally relevant complexes. In this study, we show that asukamycin, a manumycin polyketide bearing multiple electrophilic sites, reacts with C374 of the E3 ligase UBR7 to impair breast cancer cell proliferation and survival, through engaging in molecular glue interactions with multiple neo-substrates, including TP53. We demonstrate that asukamycin activates TP53 transcriptional activity in a UBR7 and C374-dependent manner and that the anti-proliferative effects observed in breast cancer cells depends on UBR7 and TP53. This molecular glue activity is also recapitulated with a related polyketide manumycin A. Furthermore, we show that manumycin D, which lacks an epoxide, still retains binding to UBR7 but shows less potent binding to TP53 and is not able to covalently modify TP53 to the same extent, consistent with this class of molecules targeting C374 of UBR7 through a hetero-Michael addition, and then engaging TP53 likely initially through reversible binding and then through a covalent bond involving the reactive epoxide. This latter observation is based on UBR7 pulldown studies showing not only the higher molecular weight TP53 species, but also the parent TP53 molecular weight. While we seem to observe one primary higher molecular weight UBR7-TP53 band, there are also several additional higher molecular weight species observed, which may correspond to multiple multi-covalent interactions with the three potential reactive sites on asukamycin and manumycin A. These complexes could also contain additional potential molecular glue partners, such as those identified in the enrichment proteomic studies. Furthermore, while we show by isoTOP-ABPP that C374 of UBR7 appears to be one of the primary targets of UBR7 and our genetic validation data shows that targeting this cysteine appears to drive the asukamycin-mediated antiproliferative effects, asukamycin and manumycin A likely possess additional targets. The previously reported targets including IKKβ, thioredoxin reductase 1, farnesyltransferase, sphingomyelinase, and LC3-mediated cytoplasmic vacuolation pathways may also play a role in mediating anti-cancer effects or other biological activities ^30,31,46–48^. Manumycin A has been shown to interfere with IKK activity by directly binding to IKKβ and forming stable high molecular mass complexes, indicating that these alternate biological activities may also be mediated through additional molecular glue-type interactions beyond those reported here ^47^.

Previous studies have demonstrated that manipulation of molecular glue scaffolds change neo-substrate scope. For example, thalidomide analogs still bind to cereblon but engage in glue interactions with different neo-substrates leading to their degradation ^19^. Rapamycin analogs glue FKBP proteins to different neo-substrate proteins leading to modulation of their function ^18,18,49^. While we showed here that both asukamycin and manumycin A stabilize the interaction between UBR7 and TP53, additional manipulation of the manumycin polyketide scaffold may result in altered neo-substrate binding profiles.

Several unanswered questions still remain. We do not understand how these manumycin polyketide/UBR7 complexes interact with p53 to activate p53 function. We also do not know the site or nature of the amino acid(s) on p53 wherein the suspected reaction with the epoxide takes place. Solving the crystal structure of the UBR7-asukamycin-TP53 ternary complex will be particularly interesting to understand the biochemical underpinnings of this molecular glue activity. We also do not understand the endogenous role of UBR7 and whether asukamycin affects this function. UBR7 belongs to the UBR family of *N*-recognin E3 ligases, but unlike UBR1-6, contains a plant homeodomain (PHD) rather than an F-box domain ^50,51^. It has been reported that UBR7 can ubiquitinylate histones and has a variety of interacting partners in other contexts ^36,52,53^. Our data currently suggest that the asukamycin-mediated phenotypes and effects upon TP53 activity are independent of ubiquitination functions of UBR7. However, we cannot exclude the normal functional role of UBR7 in conferring anti-cancer activity of asukamycin ^37,52,54^. Further work will be critical in better understanding the role of UBR7 and cancer.

Despite these remaining questions, our study showcases the possibility that multi-covalent small-molecules can potentially act as new molecular glues and that these interactions can be quickly deciphered using chemoproteomic approaches. Of future interest will be whether other natural products or synthetic small-molecules that bear multiple reactive centers can act as specific molecular glues to confer new protein functions. In addition, the ability of manumycin polyketides to template the interaction of UBR7 with multiple other proteins warrants the further examination of this natural product family for applications beyond cancer.

## Acknowledgement

We thank the members of the Nomura Research Group, the Maimone Research Group, and Novartis Institutes for BioMedical Research for critical reading of the manuscript. This work was supported by Novartis Institutes for BioMedical Research and the Novartis-Berkeley Center for Proteomics and Chemistry Technologies (NB-CPACT) for all listed authors. This work was also supported by the Nomura Research Group and the Mark Foundation for Cancer Research and Chordoma Foundation ASPIRE Award for DKN, YI. This work was also supported by grants from the National Institutes of Health (R01CA240981 for DKN, TJM, YI). YI and MO were also supported by the Japanese Society for the Promotion of Science (JSPS) postdoctoral fellowships.

## Author Contributions

YI, DKN, TJM conceived the project and wrote the paper. YI, JAT, JMK, LM, MS, TJM, DKN provided intellectual contributions and insights into project direction. YI, TJM, DKN designed the experiments. YI, MO, RW, DKN performed experiments and analyzed data. All authors edited the paper.

## Online Methods

### Materials

Asukamycin and Manumycin A were obtained from Cayman Chemicals. Heavy and light TEV-biotin tags were synthesized per previously described methods ^14^. Manumycin D synthesis and characterization is in **Synthetic Methods**. Recombinant UBR7 pure proteins were purchased from Origene or NOVUS Biologicals. Recombinant TP53 pure protein was purchased from R&D Systems.

### Cell Culture

The 231MFP cells were obtained from Prof. Benjamin Cravatt and were generated from explanted tumor xenografts of MDA-MB-231 cells as previously described ^55^. HCC38 and HEK293T cells were obtained from American Type Culture Collection (ATCC). 231MFP cells were cultured in L15 medium (HyClone) containing 10% FBS and 2 mM glutamine and maintained at 37 °C with 0% CO_2_. HEK293T cells were cultured in Dulbecco’s Modified Eagle’s Medium (DMEM) containing 10% FBS and 2 mM glutamine and maintained at 37 °C with 5% CO_2_. HCC38 cells were cultured in RPMI medium containing 10% FBS and maintained at 37 °C with 5% CO_2_.

### Survival and Proliferation Assays

Cell survival and proliferation studies were performed using Hoechst 33342 dye (Invitrogen) as described previously ^2,15,24^. Briefly, 231MFP cells were seeded at 20,000 (proliferation) or 40,000 (survival) cells/well, respectively, in serum-containing (proliferation) or serum-free (survival) media in 96-well plates and allowed to adhere overnight. The cells were treated with DMSO vehicle- or Asukamycin-containing media for 24 or 48 h before fixation and staining with 10% formalin and Hoechst 33342 dye. Studies with HCC38 cells were also performed as above but were seeded with 10,000 cells/well for proliferation and 20,000 cells/well for survival.

### IsoTOP-ABPP of Asukamycin Targets

IsoTOP-ABPP studies were done as previously reported ^2,15,24^. 231MFP cells were treated with DMSO vehicle or 10 μM of Asukamycin for 3 h. Cells were then harvested and lysed by probe sonication in PBS and protein concentrations were determined by bicinchoninic acid (BCA) assay (Pierce). Proteomes were subsequently labeled with 100μM of iodoacetamide (IA)-alkyne (CHESS Gmbh.) at room temperature (RT) for 1 h. Copper-catalyzed azide-alkyne cycloaddition (CuAAC) was performed by sequential addition of tris(2-carboxyethyl) phosphine (1 mM, Sigma), tris[(1-benzyl-1H-1,2,3-triazol-4-yl)methyl]amine (34 mM, Sigma), copper (II) sulfate (1 mM, Sigma), and biotin-linker-azide, the linker functionalized with a TEV protease recognition sequence along with an isotopically light or heavy valine for treatment of vehicle- or Asukamycin-treated proteome, respectively. After CuAAC, proteomes were precipitated by centrifugation at 6500×g, washed in ice-cold methanol, combined in a 1:1 vehicle/Asukamycin ratio, washed again, then denatured and resolubilized by heating in 1.2 % SDS/PBS at 80 °C for 5 min. Insoluble components were precipitated by centrifugation at 6500×g and soluble proteome was diluted with PBS to be 0.2% of SDS. Labeled proteins were bound to streptavidin-agarose beads (Pierce) while rotating overnight at 4 °C. After washing three times each in PBS and water, the bead-linked proteins were resuspended in 6 M urea/PBS and reduced in 1 mM of TCEP, alkylated with 18 mM of IA (Sigma), then washed and resuspended in 2 M urea and trypsinized overnight at 37 °C with 2 ug/sample sequencing grade trypsin (Promega). After washing three times each in PBS and water, the beads were resuspended in TEV buffer solution (1×TEV buffer containing 100 μM of dithiothreitol) and incubated overnight at 29 °C with Ac-TEV protease (Invitrogen). Peptides were diluted in water and acidified with 1.2 M of formic acid (Spectrum) for isoTOP-ABPP analysis. Subsequent steps of the isoTOP-ABPP and mass spectrometry analysis were performed using the same methods as we have described previously ^2,15,24^.

### Gel-Based ABPP

Gel-based ABPP methods were performed as previously described ^24^. Recombinnat pure protein (0.1μg/sample) was pre-treated with either DMSO vehicle or natural products at 37 °C for 30 min in 25 μL of PBS, and subsequently treated with 200 nM of IA-Rhodamine (Setareh Biotech) at RT for 1 h. The reaction was stopped by addition of 4×reducing Laemmli SDS sample loading buffer (Alfa Aesar). After boiling at 95°C for 5 min, the samples were separated on precast 4-20% Criterion TGX gels (Bio-Rad). Probe-labeled proteins were analyzed by in-gel fluorescence using a ChemiDoc MP (Bio-Rad).

### Constructing Knockdown Lines and Reinforced Expression or Overexpression Lines

Short-hairpin oligonucleotides were used to knock down the expression of UBR7 or TP53 in 231MFP cells using previously described methods ^2^. For lentivirus production, lentiviral plasmids and packaging plasmids (pMD2.5G, Addgene nos. 12259 and psPAX2, Addgene no. 12260) were transfected into HEK293T cells using Lipofectamine 2000 (Invitrogen). Lentivirus was collected from filtered cultured medium and used to infect the target cell line with 1:1000 dilution of polybrene. Target cells were selected over 3 days with 1 μg/ml of puromycin. The short-hairpin sequences which were used for generation of the knockdown lines were: UBR7: CCGGGATGATGTCCGGGAGGTTAAACTCGAGTTTAACCTCCCGGACATCATCTTTTTG (Sigma UBR7 MISSION shRNA Bacterial Glycerol Stock, TRCN0000294293).

TP53:CCGGCGGCGCACAGAGGAAGAGAATCTCGAGATTCTCTTCCTCTGTGCGCCGTTTTT (Sigma TP53 MISSION shRNA Bacterial Glycerol Stock, TRCN0000003753).

MISSION TRC1.5 pLKO.1- or TRC2 pLKO.5-puro Non-Mammalian shRNA Control (Sigma) was used as a control shRNA.

For expression of WT or C374A mutant UBR7, cells were transiently transfected with UBR7 expression plasmid using Lipofectamine 2000. Wild-type human UBR7 expression plasmid with C-terminal FLAG tag was purchased from Origene (RC218298). The UBR7 C374A mutant was generated with Q5 Site-Directed Mutagenesis Kit (New England Biolabs) according to manufacturer’s protocols.

### Western Blotting

FLAG antibody (M2) was obtained from Sigma. HPRT antibody (F-1) was obtained from Santa Cruz Biotechnology. Antibodies to UBR7 (PA5-31559) and p53 (DO-1) were obtained from Thermo Fisher Scientific. Antibodies to GAPDH (D16H11), Ubiquitin (P4D1), FLAG (D6W5B), and Histone H3 (D1H2) were obtained from Cell Signaling Technology.

Proteins were resolved by SDS/PAGE and transferred to nitrocellulose membranes using the iBlot system (Invitrogen). Membranes were blocked with 5 % nonfat milk or BSA in Tris-buffered saline containing Tween 20 (TBS-T) solution for 30 min at RT, washed in TBS-T, and probed with primary antibody diluted in recommended diluent per manufacturer overnight at 4 °C After 3 times washes with TBS-T, the membranes were incubated in the dark with IR800-conjugated secondary antibodies at 1:10,000 dilution in 5 % nonfat milk in TBS-T at RT for 1 h. Blots were visualized using an Odyssey Li-Cor fluorescent scanner. The membranes were stripped using ReBlot Plus Strong Antibody Stripping Solution (EMD Millipore) when additional primary antibody incubations were performed. For dual color Western blotting, anti-mouse IR680 and anti-rabbit IR800 were used to detect TP53 (DO-1) and FLAG (D6W5B), respectively.

### Anti-FLAG Pulldown

231MFP cells with stable expression of FLAG-GFP or FLAG-UBR7 were treated with DMSO vehicle or 50 μM of Asukamycin for 3 h. Cells were collected, washed twice with PBS, and lysed by probe sonication in TBS. After centrifugation at 12,000×g for 10 min, the supernatant was incubated with Anti-DYKDDDDK G1 affinity resin (GenScript) at 4 °C for 3 h. Beads were washed two times with 1 mL of cold TBS (10 min per incubation), and the samples were eluted two times by 250 μg/mL of 3×FLAG-peptide solution. Eluent was subsequently prepared for proteomic analysis as described below.

### Proteomic Analysis of Pulldown Samples

The samples were precipitated by addition of trichloroacetic acid (TCA) at a final concentration of 20% and incubated at −80°C for 1 h. The samples were then centrifuged at 14,800 rpm for 10 min and supernatant was carefully removed. After washing two times with ice cold 0.01 M HCl/90 % acetone solution, the precipitated protein was resuspended in 4 M urea containing 0.1 % Protease Max (Promega) and diluted in 40 mM ammonium bicarbonate buffer. The samples were reduced with 10 mM TCEP at 60°C for 30 min. The samples were then diluted with PBS and incubated overnight at 37 °C with sequencing grade trypsin. After centrifugation at 13,200 rpm for 30 min, the supernatant was acidified with 5 % formic acid and was subsequently analyzed by LC-MS/MS. Data was analyzed by spectral counting. Only those proteins that showed >2 peptides in at least one sample was subsequently interpreted. While this filtered and original list was interpreted for average, sem, and p-values, for visual representation of this proteomic data in **Figure 2a**, we set those proteins that showed averages <1 within a particular group as 1 to generate fold-changes that could be plotted in the figure.

### In Vitro Deubiquitination Assay

231MFP cells were treated with either DMSO vehicle or 50 μM of Asukamycin for 3 h. Cells were then harvested and lysed by probe sonication in TBS, and protein concentrations were determined by BCA assay. Cell lysates were incubated with deubiquitinating enzyme USP2 (R&D Systems) at 37 °C for 1 h. The reaction was stopped by addition of 4× reducing Laemmli SDS sample loading buffer. The reaction products were analyzed by Western blotting.

### Thermal Shift Assay

231MFP cells were lysed by probe sonication in TBS containing complete protease inhibitors cocktail (Roche), and protein concentrations were determined by BCA assay. Cell lysates (1 mg/mL) were treated with either DMSO vehicle or 50 μM of Asukamycin for 1 h at 37 °C Cell lysates were then separated into 8 fractions for thermal profiling. Fractions were heated at the indicated temperatures (42–60 °C) for 3 min using a thermal cycler (T100 Thermal Cycler, Bio-Rad), followed by 3 minutes at room temperature. Samples were then centrifuged at 16,000 × g for 15 min to separate protein aggregates from soluble proteins. Supernatants were collected and analyzed by Western blotting.

### In Vitro Transcription Factor Assay for TP53

Recombinant TP53 protein (0.05 μg/sample) was spiked into 40 μL of 231MFP breast cancer cell lysate (0.5 mg/mL in TBS) and then treated with either DMSO vehicle or 50 μM of Asukamycin for 1 h at 37 °C TP53 DNA binding to p53 DNA consensus sequence was then assessed by using the p53 transcription factor assay kit (Cayman) according to manufacturer’s protocols.

### Luciferase Reporter Assay

HEK293T cells were seeded at 30,000 cell/well in 96-well plates. After 24 h, the cells were transiently co-transfected with 0.1 μg /well of pGL4.38 [luc2P/p53 RE/Hygro] (Promega) and 0.02 μg/well of pRL-TK (renilla luciferase control vector) using Lipofectamine 2000. After treating the cells with compounds for 6 h, the expression levels of the firefly and renilla luciferase reporter genes were examined by a Dual-Glo luciferase assay system (Promega) according to the manufacturer’s protocols. The luminescent signals were measured using a SpectraMax i3 plate reader (Molecular Devices).

### Data Availability Statement

The datasets generated during and/or analyzed during the current study are available from the corresponding author on reasonable request.

## Synthetic Methods

### General Procedures

Unless otherwise stated, all reactions were performed in oven-dried or flame-dried Fisherbrand® borosilicate glass tubes (Fisher Scientific, 1495925A, 13 × 100 mm) with a black phenolic screw cap (13-425) under an atmosphere of dry nitrogen. Manumycin A was purchased from Cayman Chemicals, and used directly without further purification. Reactions were monitored by thin layer chromatography (TLC) on TLC silica gel 60 F_254_ glass plates (EMD Millipore) and visualized by UV irradiation and staining with *p*-anisaldehyde, phosphomolybdic acid, or potassium permanganate. Volatile solvents were removed under reduced pressure using a rotary evaporator. Flash column chromatography was performed using Silicycle F60 silica gel (60Å, 230-400 mesh, 40-63 μm). Ethyl acetate and hexanes were purchased from Fisher Chemical and used for chromatography without further purification. Proton nuclear magnetic resonance (^1^H NMR) and carbon nuclear magnetic resonance (^13^C NMR) spectra were recorded on a Bruker AV-600 spectrometer operating at 600 MHz for ^1^H, and 150 MHz for ^13^C. Chemical shifts are reported in parts per million (ppm) with respect to the residual solvent signal CDCl_3_ (^1^H NMR: δ = 7.26; ^13^C NMR: δ = 77.16). Peak multiplicities are reported as follows: *s* = singlet, *d* = doublet, *t* = triplet, *dd* = doublet of doublets, *tt* = triplet of triplets, *m* = multiplet, *br* = broad signal, *app* = apparent. High-resolution mass spectra (HRMS) were obtained by the QB3/chemistry mass spectrometry facility at the University of California, Berkeley.

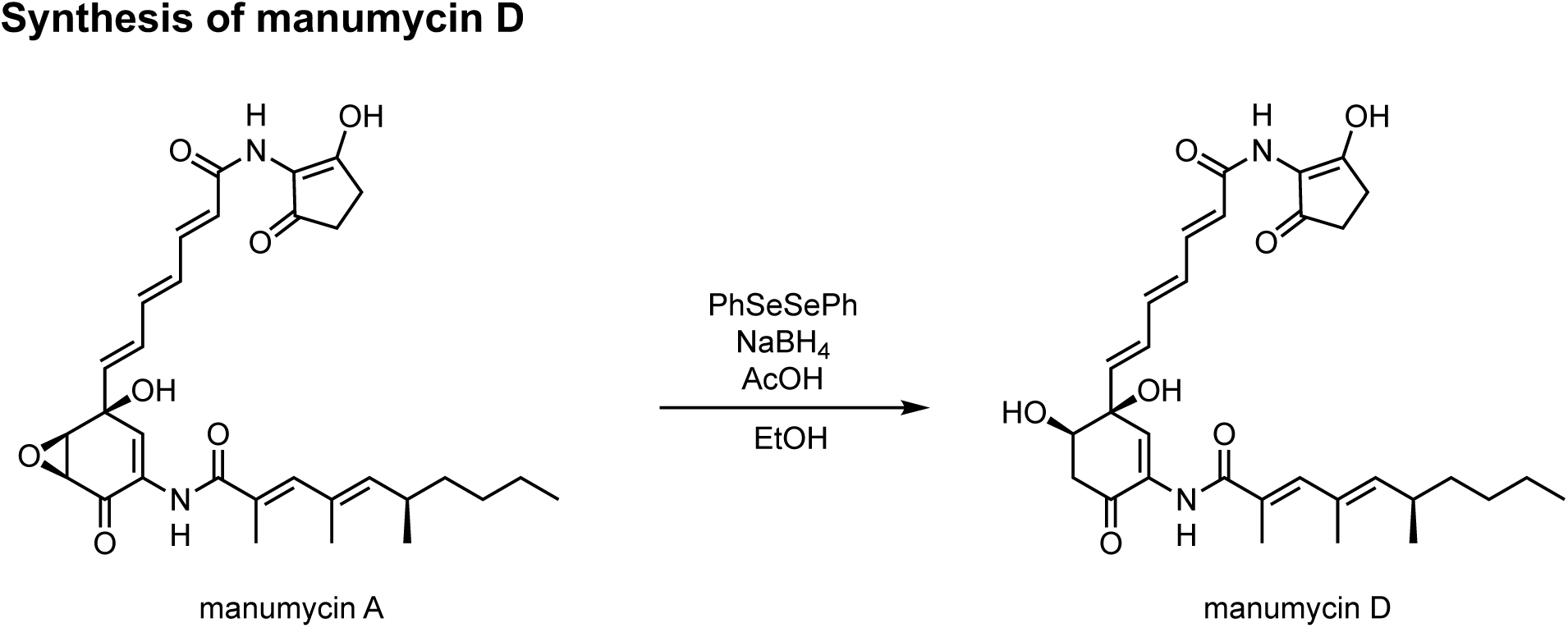

### Synthesis of manumycin D

The following procedure was conducted based on a slight modification to known procedures ^1,2^. To a stirring solution of diphenyl diselenide (4.2 mg, 0.015 mmol, 1.5 eq.) in ethanol (0.15 mL) under a nitrogen atmosphere was added sodium borohydride (1.0 mg, 0.027 mmol, 3.0 eq.) at room temperature. After evolution of hydrogen ceased, the light-yellow solution was cooled to 0 °C, followed by the addition of acetic acid (0.52 µL, 0.015 mmol, 1.5 eq.), and the mixture was further stirred at 0 °C for 5 min. The resulting light-yellow solution was then added to a solution of manumycin A (5.0 mg, 0.0091 mmol, 1.0 eq.) in ethanol (0.1 mL) at room temperature, and upon complete consumption of manumycin A (monitored by TLC, 5 min) the reaction mixture was diluted with ethyl acetate (1.0 mL), and oxygen was passed through the solution for 5 minutes. Volatiles were removed under reduced pressure, and dichloromethane (1.0 mL) was added. Insoluble materials were removed by passing the mixture through a plug of celite, and the crude material was purified by preparative TLC (CHCl_3_:MeOH = 9:1) to give manumycin D (2.9 mg, 58% yield) as a dark-orange oil whose data was in agreement with that previously reported ^3,4^: *R*f = 0.33 (SiO_2_, CHCl_3_:MeOH = 9:1, UV, KMnO_4_); ^1^H NMR: (600 MHz, CDCl_3_) δ 13.54 (br, 1H), 8.29 (br, 1H), 7.56 (m, 2H), 7.32 (dd, *J* = 14.8, 11.3 Hz, 1H), 6.80 (s, 1H), 6.66 – 6.51 (m, 2H), 6.46 – 6.36 (m, 1H), 6.09 – 5.97 (m, 2H), 5.34 (d, *J* = 9.6 Hz, 1H), 4.17 – 4.03 (m, 1H), 3.23 (br, 1H), 2.90 (dd, *J* = 17.1, 6.1 Hz, 1H), 2.84 (br, 1H), 2.76 (dd, *J* = 17.1, 3.4 Hz, 1H), 2.65 – 2.58 (m, 2H), 2.55 – 2.50 (m, 2H), 2.50 – 2.39 (m, 1H), 2.06 (s, 3H), 1.82 (s, 3H), 1.39 – 1.22 (m, 6H), 0.98 (d, *J* = 6.6 Hz, 3H), 0.89 (t, *J* = 7.0 Hz, 3H); ^13^C NMR (150 MHz, CDCl_3_) δ 197.4, 191.9, 174.0, 169.0, 165.5, 143.7, 142.9, 140.3, 139.9, 138.7, 132.4, 131.6, 131.5, 130.1, 128.5, 126.1, 121.5, 115.1, 73.7, 71.9, 40.7, 37.2, 33.0, 32.3, 29.9, 25.8, 23.0, 20.9, 16.7, 14.23, 14.18; HRMS: (ESI-TOF, m/z) *calcd*. For C_31_H_39_N_2_O_7_ [M–H]^−^ calc.: 551.2763; found: 551.2765.

