## Supplementary Information for "Manumycin Polyketides Act as Molecular Glues Between UBR7 and P53 to Impair Breast Cancer Pathogenicity"

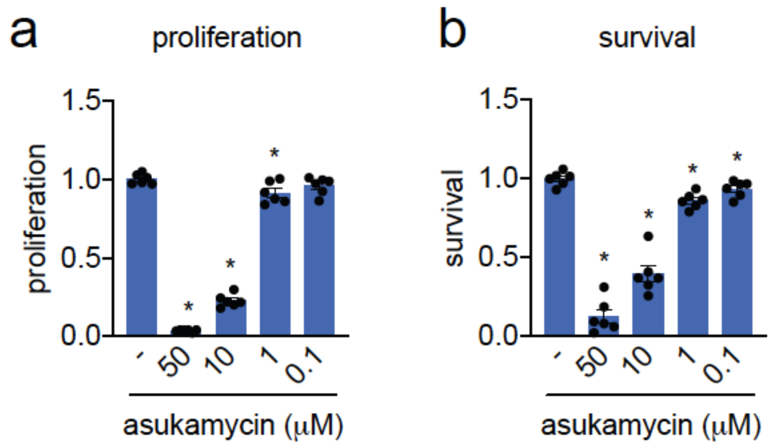


**Supplementary Figure 1. Asukamycin effects in HCC38 cells.** Proliferation and serum-free cell survival in HCC38 breast cancer cells treated with DMSO vehicle or asukamycin for 48 h, assessed by Hoechst stain. Data shown as individual replicate values and average ± sem and are n=6 biologically independent samples/group. Statistical significance was calculated with two-tailed unpaired Student's t-tests and are shown as *$p<0.05$ compared to vehicle-treated controls within each group.

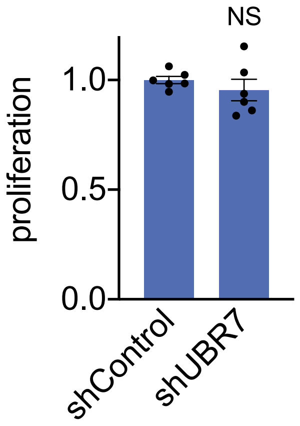


**Supplementary Figure 2. Proliferation in 231MFP cells.** Proliferation in 231MFP shControl and shUBR7 breast cancer cells for 48 h, assessed by Hoechst stain. Data shown as individual replicate values and average ± sem and are n=6 biologically independent samples/group. NS denotes that shUBR7 proliferation is not significantly changed ($p>0.05$) compared to the shControl group by a Student's two-tailed t-test.

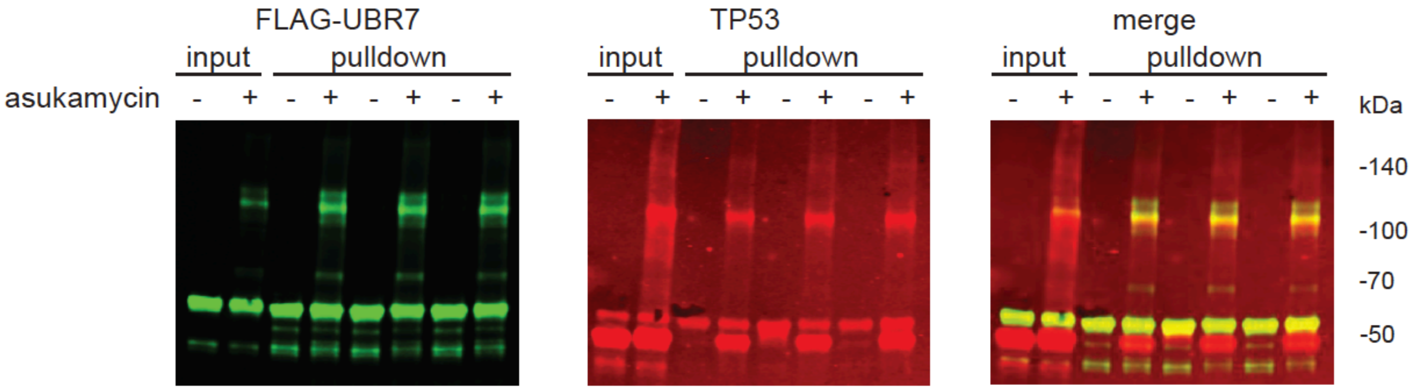


**Supplementary Figure 3. Dual color Western blots of FLAG-UBR7**. 231MFP cells stably expressing FLAG-GFP or FLAG-UBR7 were treated with DMSO vehicle or asukamycin (50 μM) for 3 h. FLAG-GFP and FLAG-UBR7 interacting proteins were subsequently enriched and then subjected to dual color Western blotting analysis of FLAG-UBR7 (in green) and TP53 (in red). Blot shown on the right is a merged blot of FLAG-UBR7 and TP53 with the yellow color indicating overlap.

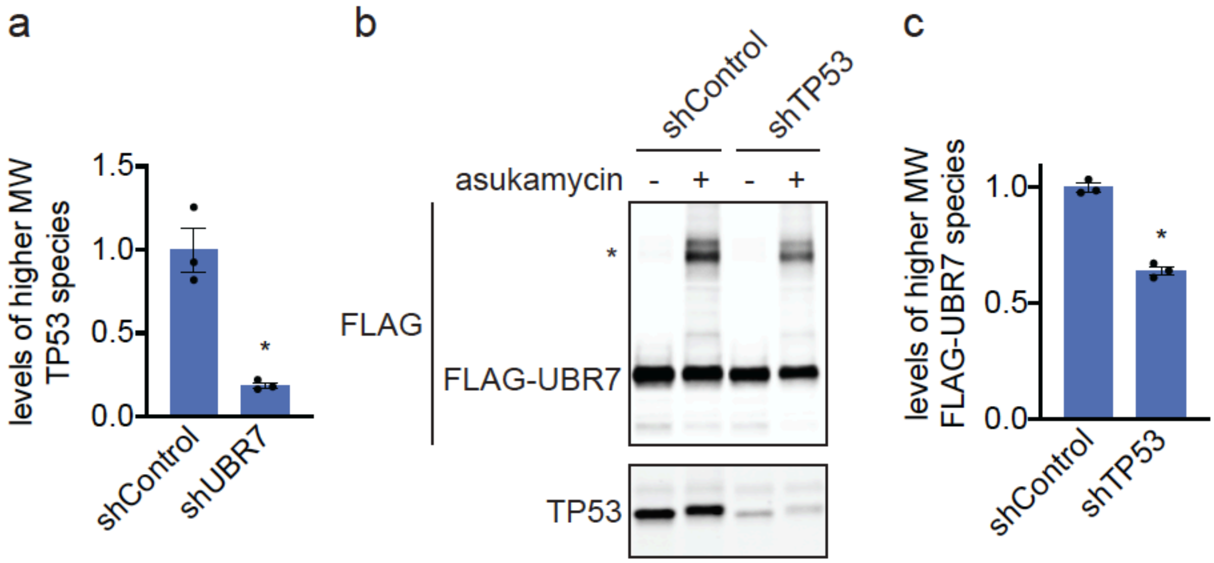


**Supplementary Figure 4. Understanding the composition of the asukamycin-mediated higher molecular weight band. (a)** Quantification of TP53 levels normalized to GAPDH levels in Western blot shown in **Figure 2d. (b)** Anti-FLAG and anti-TP53 blot in shControl and shTP53 231MFP breast cancer cells expressing FLAG-UBR7 treated with vehicle DMSO or asukamycin (50 μM) for 3 h. * notes the higher molecular FLAG-UBR7 band. **(c)** Quantification of higher molecular weight FLAG-UBR7 band noted with * in **(b)**. Data shown in **(a**, **c)** as individual replicate values and average ± sem and are n=3 biologically independent samples/group. Gel shown in **(b)** is a representative gel of n=3 biologically independent samples/group. Statistical significance was calculated with two-tailed unpaired Student's t-tests and are shown as *p<0.05 compared to shControl cells treated with asukamycin in **(a, c)**.

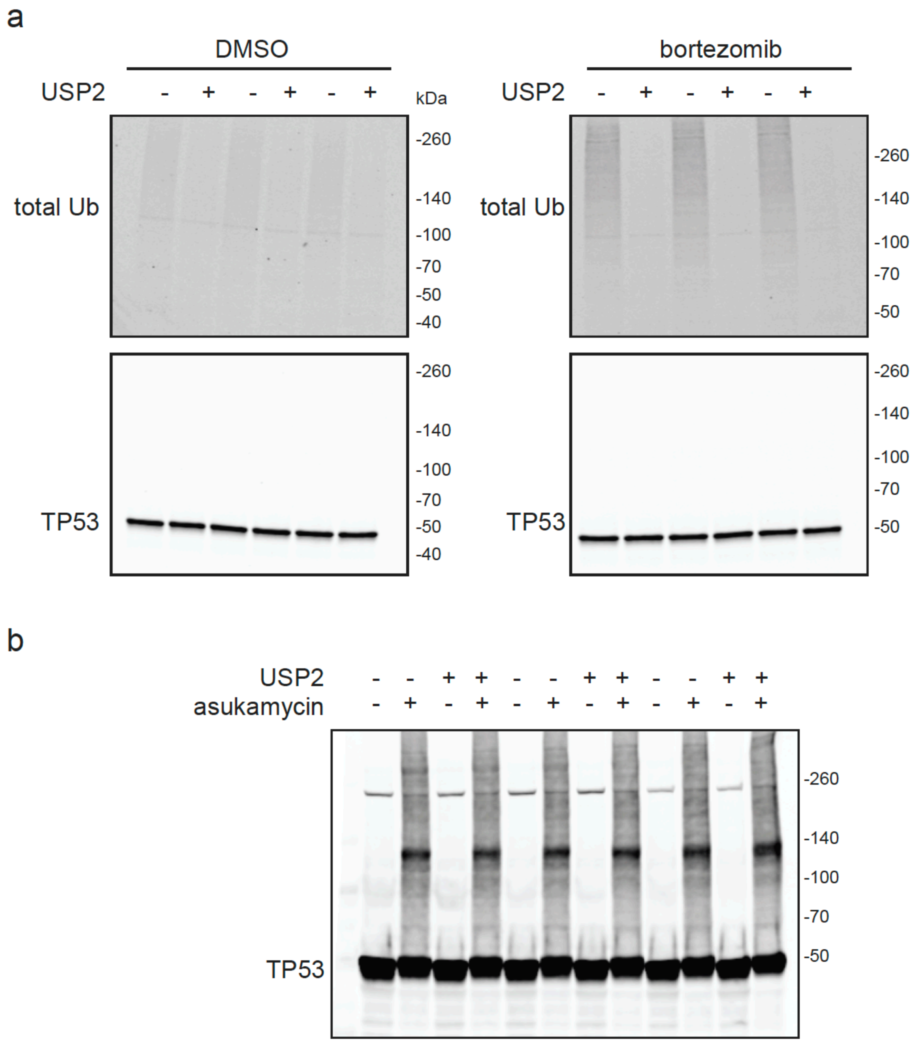


**Supplementary Figure 5. Asukamycin-mediated higher molecular weight TP53 species are not polyubiquitinated protein. (a)** Total proteome ubiquitin Western blot from 231MFP cell lysate treated with deubiquitinase USP2 (0.3 micrograms, 1 h). 231MFP cells were treated either with DMSO vehicle or proteasome inhibitor bortezomib (1 μM, 3 h) to accumulate ubiquitinated proteins. This experiment is a positive control experiment to show that our conditions for USP2 incubation can show removal of ubiquitin from polyubiquitinated proteins. **(b)** 231MFP cells treated *in situ* with DMSO vehicle or asukamycin (50 μM) for 4 h. Cell lysates were then treated with USP2 (0.3 micrograms, 1 h). The 260 kDa band is likely a non-specific band not corresponding to TP53.

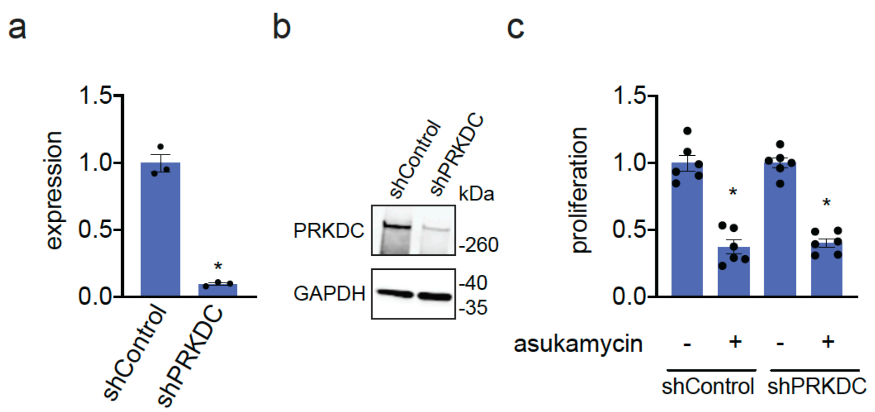


**Supplementary Figure 6.** PRKDC knockdown in 231MFP breast cancer cells. **(a, b)** PRKDC was stably knocked down using shRNAs and PRKDC knockdown was confirmed by qPCR **(a)** and by Western blotting **(b)** compared to shControl cells**. (c)** Proliferation in 231MFP breast cancer cells treated with DMSO vehicle or asukamycin (50 μM, 48 h), and assessed by Hoechst staining. Data shown as individual replicate values and average ± sem and are n=3 biologically independent samples/group for **(a, b)** and n=6 biologically independent samples/group for **(c)**. Statistical significance was calculated with two-tailed unpaired Student's t-tests and are shown as *p<0.05 compared to shControl groups in **(a, b)** and compared to respective shControl or shPRKDC vehicle-treated controls in **(c)**.

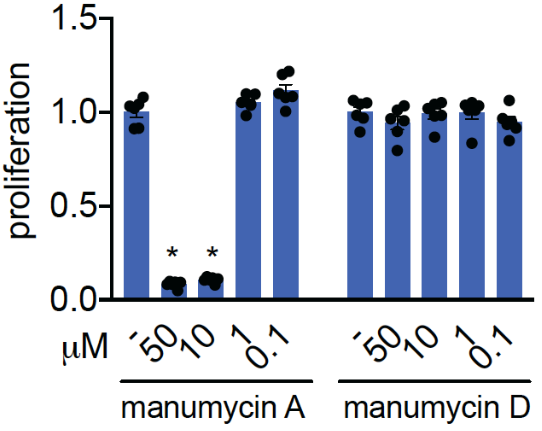


**Supplementary Figure 7.** Proliferation in 231MFP breast cancer cells treated with manumycin A or manumycin D for 48 h and assessed by Hoechst staining. Data shown as individual replicate values and average ± sem and are n=6 biologically independent samples/group. Statistical significance was calculated with two-tailed unpaired Student's t-tests and are shown as *p<0.05 compared to vehicle-treated controls within each group.

**Supplementary Datasets**

**Supplementary Dataset 1. IsoTOP-ABPP analysis of asukamycin treatment *in situ* in 231MFP breast cancer cells.** IsoTOP-ABPP analysis of asukamycin treatment *in situ* (10 μM). 231MFP breast cancer cells were treated with DMSO or asukamycin (10 μM, 3 h *in situ*), after which cells were harvested and proteomes were labeled *ex situ* with IA-alkyne (100 μM, 1 h), followed by appendage of isotopically light (for DMSO-treated) or heavy (for asukamycin-treated) TEV protease cleavable biotin-azide tags by copper-catalyzed azide-alkyne cycloaddition (CuAAC). Control and treated proteomes were subsequently combined in a 1:1 ratio, probe-labeled proteins were avidin-enriched, digested with trypsin, and probe-modified tryptic peptides were eluted by TEV protease, analyzed by LC-MS/MS, and light to heavy probe-modified peptide ratios were quantified. Shown are data from n=3 biological replicates/group.
**Tab 1.** Total isoTOP-ABPP proteomic dataset
**Tab 2.** Analyzed isoTOP-ABPP dataset. For those probe-modified peptides that showed ratios >2, we only interpreted those targets that were present across all three biological replicates, were statistically significant, and showed good quality MS1 peak shapes across all biological replicates. Light versus heavy isotopic probe-modified peptide ratios are calculated by taking the mean of the ratios of each replicate paired light vs. heavy precursor abundance for all peptide spectral matches (PSM) associated with a peptide. The paired abundances were also used to calculate a paired sample t-test p-value in an effort to estimate constancy within paired abundances and significance in change between treatment and control. P-values were corrected using the Benjamini/Hochberg method.

**Supplementary Dataset 2**. **Proteomics analysis of molecular glue interactions with UBR7-asukamycin.** 231MFP cells stably expressing FLAG-GFP or FLAG-UBR7 were treated with DMSO vehicle or asukamycin (50 μM) for 3 h. FLAG-GFP and FLAG-UBR7 interacting proteins were subsequently enriched and then subjected to proteomic analysis. Data were quantified by spectral counting. Raw data are shown in Tabs 1 and 2. Tab 3 shows data for those proteins that showed at least 2 spectral counts in at least one sample. For those proteins that showed no peptides in a particular group, we set those proteins to 1 to enable relative fold-change quantification to generate **Figure 2a** to visually show those proteins that showed high levels in asukamycin (ASK)-treated FLAG-UBR7 groups compared to DMSO-treated FLAG-GFP groups on the x-axis and ASK-treated FLAG-UBR7 groups compared to DMSO-treated FLAG-UBR7 groups. These data are shown in Tabs 4 and 5. The hits that we designated as potential hits for molecular glue-like interactions with UBR7 and asukamycin were those proteins that showed >15-fold comparing UBR-ASK to UBR7-DMSO groups and >4-fold enrichment comparing UBR7-ASK to GFP-ASK groups.
